# A Bayesian computational approach to explore the optimal duration of a cell proliferation assay

**DOI:** 10.1101/147678

**Authors:** Alexander P Browning, Scott W McCue, Matthew J Simpson

## Abstract

Cell proliferation assays are routinely used to explore how a low density monolayer of cells grows with time. For a typical cell line with a doubling time of 12 hours (or longer), a standard cell proliferation assay conducted over 24 hours provides excellent information about the low-density exponential growth rate, but limited information about crowding effects that occur at higher densities. To explore how we can best detect and quantify crowding effects, we present a suite of in silico proliferation assays where cells proliferate according to a generalised logistic growth model. Using approximate Bayesian computation we show that data from a standard cell proliferation assay cannot reliably distinguish between classical logistic growth and more general non-logistic growth models. We then explore, and quantify, the trade-off between increasing the duration of the experiment and the associated decrease in uncertainty in the crowding mechanism.

## 1 Introduction

Two-dimensional *in vitro* cell biology experiments play an invaluable role in improving our understanding of the collective behaviour of cell populations (Laing, 2007). Understanding collective cell behaviour is relevant to a number of normal and pathological processes, such as tissue regeneration and malignant spreading, respectively. One of the most common *in vitro* cell biology experiments is called a *proliferation assay* (Bosco et al., 2015; Bourseguin et al., 2016). Cell proliferation assays are initiated by uniformly placing a monolayer of cells, at low density, on a two-dimensional substrate. Individual cells in the population undergo both movement and proliferation events, and the assay is observed as the density of the monolayer of cells increases. Comparing cell proliferation assays with and without a putative drug plays an important role in drug design (Bosco et al., 2015; Bourseguin et al., 2016).

One approach to interpret a cell proliferation assay is to use a mathematical model. This approach can provide quantitative insight into the mechanisms involved (Maini et al., 2004; Sengers et al., 2007). For example, it is possible to estimate the proliferation rate of cells by calibrating a mathematical model to data from a cell proliferation assay. Results can then be used to compare a target and control assay (Johnston et al., 2015). Typically, most previous studies that interpret cell biology assays using continuum mathematical models make the assumption that cells proliferate logistically (Cai et al. 2007; Dale et al., 1994; Doran et al., 2009; Jin et al., 2016a; Maini et al., 2004a; Maini et al., 2004b; O’Dea et al., 2012; Savla et al., 2004; Sengers et al., 2007; Sheardown and Cheng, 1996; Sherratt and Murray, 1990). The classical logistic equation is given by

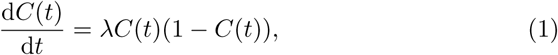

where *C*(*t*) is the scaled cell density, such that *C*(*t*) = 1 represents the carrying capacity density, *t* is time and *λ* is the cell proliferation rate. For example, by calibrating the solution of Eq (1) to data from a cell biology assay, Treloar et al. (2014) showed that the proliferation rate of 3T3 fibroblast cells is approximately 0.048 /hour. However, while the classical logistic model is routinely used to study biological population dynamics (Pearl, 1927; Edelstein-Keshet, 1988; Murray, 2002), this choice is often made without a careful examination of whether the classical logistic model is valid (Treloar et al., 2014).

In the literature, there is an awareness that biological populations do not always grow according to the classical logistic equation (Gerlee, 2013; Zwietering et al., 1990). For example, West and coworkers investigate the growth of cell populations from a wide range of animal models and find that the growth is not logistic; instead, they find that a more general model provides a better match to the experimental data (West et al., 2001). Likewise, Laird (1964) examines tumour growth data and shows that the Gompertz growth law matches the data better than the classical logistic model. Similar observations have also been made more recently for different types of tumour growth by Sarapata and de Pillis (2014).

Therefore, it is not always clear that the classical logistic model ought to be used to describe cell proliferation assays. The classical logistic model, and its generalisations (Tsoularis and Wallace, 2002), all lead to similar growth dynamics during the early phase of the experiment when the density is small. The key differences between these models occur at larger densities as the cell population grows towards the carrying capacity density. The question of whether cells in a proliferation assay grow logistically, or by some other mechanism, is obscured by the fact that most cell proliferation assays are conducted for a relatively short period of time. To illustrate this, we note that a typical cell proliferation rate of *λ* = 0.048 /hour (Treloar et al., 2014) corresponds to a doubling time of approximately 14 hours. Given that a typical initial cell density in a cell proliferation assay is approximately *C*(0) ≈ 0.1, and the typical time scale of a cell proliferation assay is no more than 24 hours, the cell density will grow to be no more than 0.4, Fig 1(a)-(d). Indeed, the evolution of the cell density data in Fig 1(d) shows that the cell density grows approximately linearly over the standard experimental duration of 24 hours. This linear increase is consistent with the early time behaviour of the exponential growth phase, but provides less information about later time behaviour where crowding effects play a role. Therefore, standard experimental durations are inappropriate for the purposes of examining how cells grow at high densities. The focus of the current work is to explore how we can determine the optimal duration of a cell proliferation assay so that it can be used to reliably distinguish between classical logistic and generalised logistic growth models. In summary, this study is the first time that an individual based model has been used to explore the duration of a cell proliferation assay, in order to reliably distinguish between different types of growth models.

**Fig. 1.**
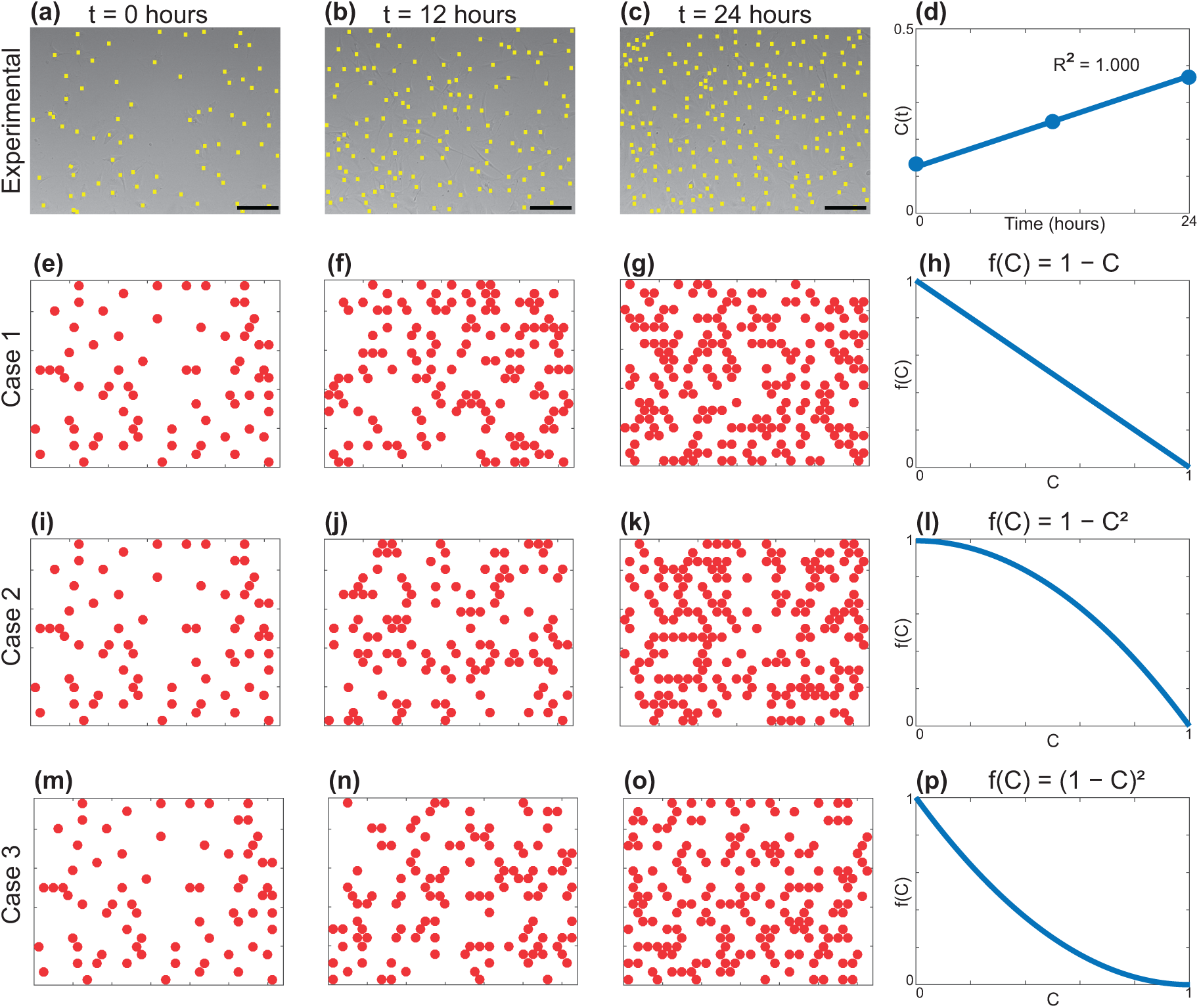
Snapshots of experimental and in *silico* cell proliferation assays. (a)-(c) show experimental images obtained from a typical cell proliferation assay performed using 3T3 fibroblast cells, shown at 12 hour intervals, as indicated. The position of each cell is identified with a yellow marker. Each scale bar corresponds to 100 *μ*m. (d) shows the scaled cell density from this experiment at each time point, and a least squares linear fit. (e),(i) and (m) show the initial distribution of agents in the in *silico* assays. The location of agents in (e),(i) and (m) are identical, and chosen so that the location of each cell in (a) is mapped to a hexagonal lattice, and placed on the nearest vacant lattice site. Simulation results are shown for: Case 1 in (e)-(g); Case 2 in (i)-(k); and, Case 3 in (m)-(o). The crowding function used in each Case is shown (h),(l) and (p) for Cases 1, 2 and 3 respectively. All simulations correspond to *I* = 25, *J* = 22, *P_m_* = 0.579, *P_P_* = 0.002, *τ* = 0.0417 hour and *Δ* = 25 pm. These discrete parameters correspond to *λ* = 0.048 /hour and *D* = 2200 *μ*m^2^/hour. Images in (a)-(c) are reprinted with permission from the Bulletin of Mathematical Biology (2013) Simpson et al. Experimental and modelling investigation of monolayer development with clustering. 75, 871-889.

This work is organised as follows. We first present a suite of results from a stochastic *in silico* cell proliferation assay. The benefit of working with an *in silico* assay is that it can be used to describe the evolution of a cell proliferation assay corresponding to a known, but general, proliferation mechanism,

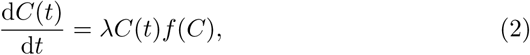

where *f*(*C*) ∈ [0, 1] is a crowding function of our choice (Jin et al., 2016b). The crowding function is a smooth decreasing function that satisfies *f*(0) = 1 and *f*(1) = 0. In general, we could study any choice of *f*(*C*) that satisfies these conditions. However, for the purposes of this study we restrict our attention to the family of crowding functions given by

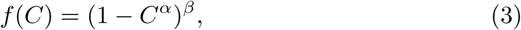

where *α* and *β* are positive constants (Tsoularis and Wallace, 2002). This choice of *f*(*C*) is still general and we note that different choices of *α* and *β* correspond to well-known biological growth models such as the classical logistic growth model, the Gompertz growth model, and the von Bertalanaffy growth model (Tsoularis and Wallace, 2002). Our choice of *f*(*C*) is partly motivated by the recent work of Sarapata and de Pillis (2014), who explore a range of sigmoid growth models for different types of tumours, including bladder, breast, liver, lung, and melanoma tumours. Sarapata and de Pillis (2014) show that the classical logistic growth model does not always provide the best match to observed data, and they test a range of other sigmoid growth models for each different kind of tumour. The different forms of sigmoid growth models that Sarapata and de Pillis (2014) explore are encompassed in our choice of crowding function, Eq 3, simply by making different choices of the constants *α* and *β*.

In this work we focus on three particular choices of *f*(*C*):

Case 1: *α* = 1 and *β* = 1. Here, *f*(*C*) is a linear function that corresponds to the classical logistic equation (Eq 1). See Fig 1(h);

Case 2: *α* = 2 and *β* = 1. Here, *f*(*C*) is a non-linear, concave-down function. See Fig 1(l); and,

Case 3: *α* = 1 and *β* = 2. Here, *f*(*C*) is a non-linear, concave-up function. See Fig 1(p).

Setting *α* = 1 and *β* = 1 recovers the classical logistic equation (Eq 1), whereas other choices of *α* and *β* lead to different, general logistic growth models. Typical *in silico* experiments showing snapshots of the growing populations are given in Fig 1(e)-(g) for Case 1, Fig 1(i)-(k) for Case 2 and Fig 1(m)-(o) for Case 3. After we have generated typical *in silico* results for these different choices of *f*(*C*), we then examine our ability to distinguish between data corresponding to different choices of *f*(*C*) using approximate Bayesian computation (ABC) (Liepe et al. 2014; Sunnaker et al. 2013; Tanaka et al. 2006; Collis et al. 2017) to estimate the parameters *α* and *β*. This procedure clearly shows that the duration of a standard cell proliferation assay is too short to reliably recover the values of *α* and *β*. Therefore, to provide quantitative insight into the benefit of performing the experiment for a longer duration, we quantify the decrease in our uncertainty of the parameters and the increase in information as we effectively run the experiment for longer periods of time.

## 2 Methods

### 2.1 Discrete mathematical model

We use a lattice-based random walk model to describe a cell proliferation assay (Liggett, 1999). Throughout the work, we will refer to a realisation of the stochastic model as either an *in silico* experiment, or a simulation. In the model cells are treated as equally-sized discs, and this is a typical assumption (Deroulers et al. 2009; Vo et al. 2015) that is supported by experimental measurements (Simpson et al. 2013). We use a hexagonal lattice, with no more than one agent per site. The lattice spacing, Δ, is chosen to be equal to the mean cell diameter (Jin et al., 2016b). This means we have a circular packing of agents, which corresponds to the maximum carrying capacity for a population of uniformly sized discs. The relationship between the scaled density, *C*(*t*), and the number of agents, *N*(*t*), is

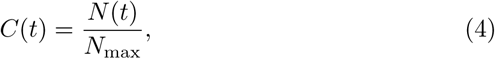

so that *C*(*t*) = 1 corresponds to the carrying capacity of *N*_max_ agents, which is the number of lattice sites. Motivated by the experimental images of the cell proliferation assay in Fig 1(a)-(c), that is conducted with 3T3 fibroblast cells, we set Δ = 25 *μ*m to be the mean cell diameter (Simpson et al., 2013). As the images in Fig 1(a)-(c) show a fixed field of view that is much smaller than the spatial extent of the uniformly distributed cells in the experiment, we apply zero net flux boundary conditions (Johnston et al., 2015).

Each lattice site, indexed (*i*, *j*) where *i*, 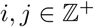, has position

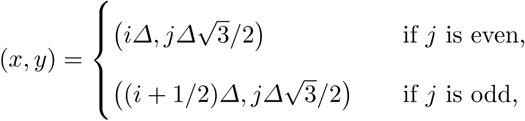

such that 1 ≤ *i* ≤ I and 1 ≤ *j* ≤ *J*. To match a typical physical domain, such as the experiment in Fig 1(a)-(c) where the field of view is 625 *μ*m × 480 *μ*m and the cell diameter is *Δ* = 25 *μ*m, we set *I* = 25 and *J* = 22. When this domain is packed to confluence, the field of view can hold no more than *N*_max_ = 550 agents.

In any single realisation of the discrete model, the occupancy of site s is denoted *C***_s_**, with *C***_s_** = 1 if the site is occupied, and *C***_s_** = 0 if vacant. We report results from the model by summing the total number of agents at time t, which we denote *N*(*t*). Each site **s** is associated with a unique index (*i, j*). We denote the set of nearest neighbour sites surrounding site s as 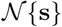, and the size of 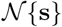 is 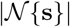. For a typical lattice site, not on any boundary, 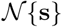 corresponds to the usual six nearest neighbour sites and 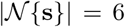. However, for any lattice site on a boundary, we adjust 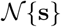 and 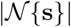 as appropriate to enforce no-flux boundary conditions.

To initiate simulations of a cell proliferation assay, we randomly select a lattice site and place an agent on that site, provided the site is vacant. We repeat this process until *N*(0) = 55 agents have been randomly placed. This corresponds to each simulation starting with *C*(0) = 0.1, which is typical of the initial density, such as in Fig 1(a). The following algorithm is used to simulate the way in which cells migrate and proliferate during the experiment. At any time, *t*, there are *N*(*t*) agents on the lattice. In each discrete time step, of duration *τ*, we allow motility and proliferation events to occur in the following two sequential steps.

First, *N*(*t*) agents are selected independently at random, one at a time with replacement, and given the opportunity to move with probability *P_m_* ∈ [0, 1]. A motile agent attempts to move to one of the six nearest neighbour sites, selected at random. To simulate crowding effects, potential motility events are aborted if an agent attempts to move to an occupied site or attempts to move outside the domain.

Second, another *N*(*t*) agents are selected independently, at random, one at a time with replacement, and given the opportunity to proliferate with probability *P_p_* ∈ [0, 1]. To assess how crowding affects the ability of a cell to proliferate, we follow the approach of Jin et al. (2016b) and assume that an agent at site s senses the occupancy of the six nearest neighbour sites, and can detect a measure of the average occupancy of those sites,

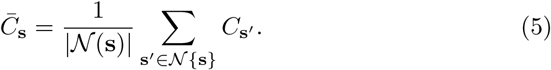

This means that 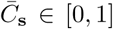 is a measure of the local crowdedness in 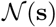. We use 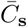 to determine whether a potential proliferation event succeeds by introducing a *crowding function*, *f*(*C*) ∈ [0, 1] with *f*(0) = 1 and *f*(1) = 0. To incorporate crowding effects we sample a random number, *R* ~ *U*(0,1). If 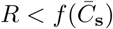, a daughter agent is placed at a randomly chosen, vacant, nearest neighbouring site, whereas if 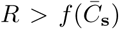, the potential proliferation event is aborted. After the *N*(*t*) potential proliferation events have been attempted, *N*(*t* + *τ*) is updated.

These two steps are repeated until the desired end time, *T*, is reached.

As previously demonstrated (Jin et al., 2016b), the continuum limit description of this discrete model gives rise to
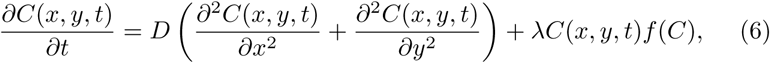

where,
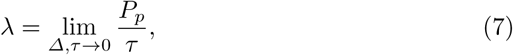

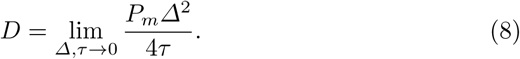

Here, *λ* is the proliferation rate, and the motility of agents is characterised by a diffusivity, *D*. Since the agents are initially distributed uniformly we have ∂*C*(*x*, *y*, *t*)*/*∂*x* ≈ *∂C* (*x*, *y*, *t*)/∂*y* ≈ 0. This means that the partial differential equation simplifies to an ordinary differential equation,

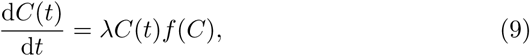

which is a generalised logistic growth model.

In this study, we only ever vary the parameters in the crowding function, *α* and *β*. All other parameters are fixed, and chosen to represent a typical cell population. As previously stated, we set *N*(0) = 55, *I* = 25 and *J* = 22, to accommodate the typical geometry and initial condition of a cell proliferation assay with a population of cells whose mean diameter is *Δ* = 25 *μ*m (Simpson et al., 2013). To describe the rate at which cells move, we set *P_m_* = 0.579 and *τ* = 0.0417 hours. This corresponds to *D* = 2200 *μ*m^2^/hour, which is a typical value of the cell diffusivity for a mesenchymal cell line (Simpson et al., 2014). To describe the rate at which cells proliferate, we set *P_p_* = 0.002 and *τ* = 0.0417 hours. This corresponds to **λ** = 0.048 /hour, which is a typical value of the cell proliferation rate (Treloar et al., 2014). This proliferation rate is consistent with the experimental data in Fig 1(d).

Using these parameter estimates, we show the evolution of *C*(*t*) for a single realisation of the discrete model, for each choice of crowding function, in Fig 2(a)-(b), for *T* = 24 and 96 hours, respectively. Results in Fig 2(a)-(b) show some stochastic fluctuations, as expected. To approximate the expected behaviour, we perform 20 identically prepared realisations of the discrete model and show the mean density profile, 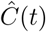, in Fig 2(c)-(d), for *T* = 24 and 96 hours, respectively. Comparing the single realisations with the mean behaviour confirms that there are minimal fluctuations, at this scale. Furthermore, we see minimal differences in the overall behaviour of the model when we consider a single realisation and the results from an ensemble of 20 identically prepared realisations.

**Fig. 2.**
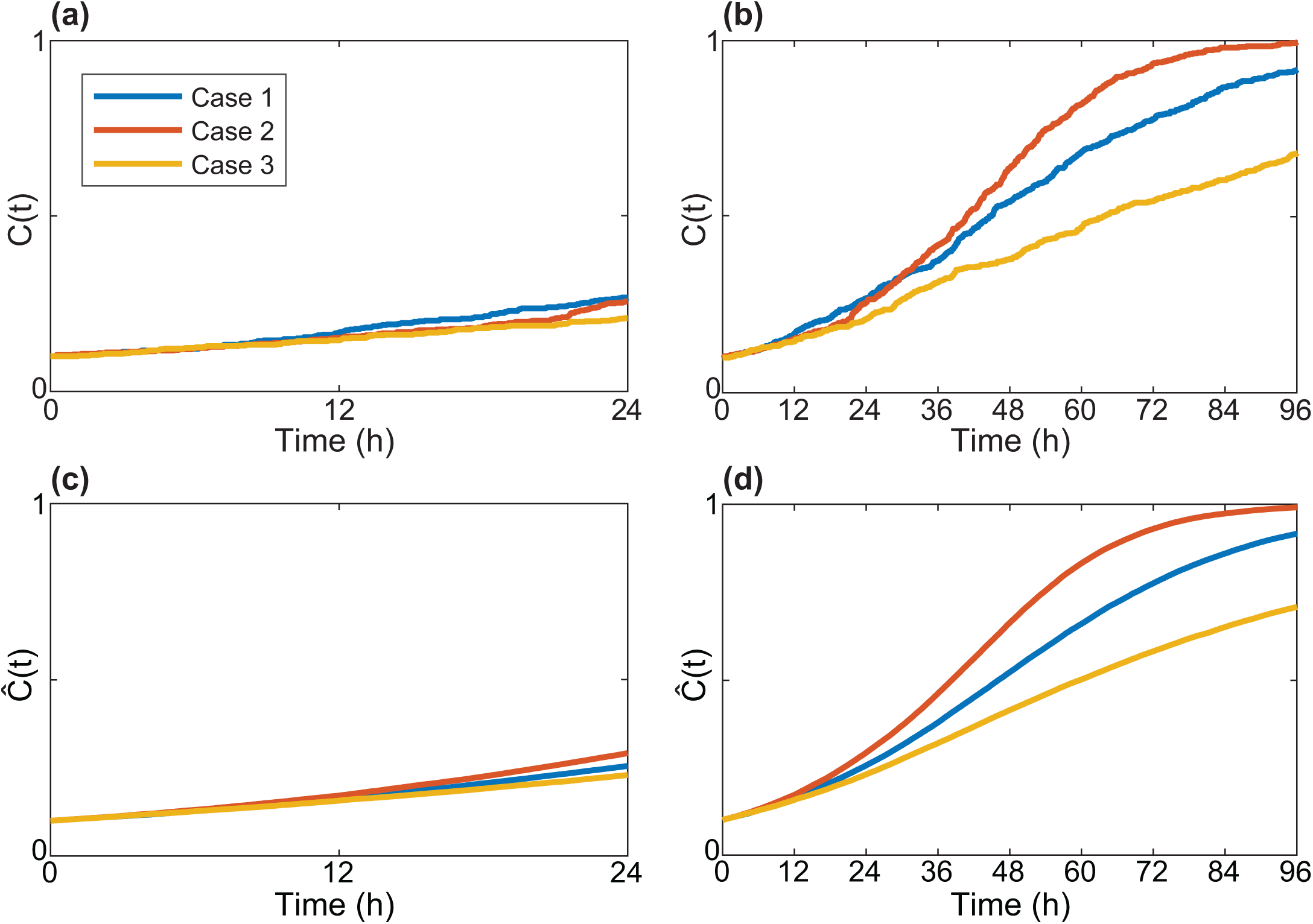
The role of the crowding function and timing for a typical proliferation assay. Results show the evolution of: (a) *C(t),* for *T* = 24 hours; (b) *C(t),* for *T* = 96 hours; (c) 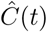, for *T* = 24 hours; and, (d) 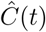, for *T* = 96 hours. All simulations correspond to *N*(0) = 55, *I* = 25, *J* = 22, *P_m_* = 0.579, *P_p_* = 0.002, *τ* = 0.0417 hours and *Δ* = 25 *μ*m. These discrete parameters correspond to *λ* = 0.048 /hour and *D* = 2200 *μ*m^2^/hour. The carrying capacity is *N_max_* = 550.

### 2.2 Parameter estimation using ABC rejection

Using a Bayesian framework, we consider the crowding function parameters ***θ*** = (*α, β*) as random variables, and the uncertainty in the ***θ*** is updated using observed data (Gelman et al., 2004; Tanaka et al., 2006; Sunnaker et al., 2013; Collis et al., 2017). Under this assumption, we note that the cell density profile, *C*(*t*), is also a random variable. In this section we refer to the variables using vector notation to keep the description of the inference algorithm as succinct as possible. However, in the main text we refer to the variables using ordered pairs, (*α*, *β*), so that our results are presented as clearly as possible.

To begin with, we perform three *in silico* experiments with fixed, known parameter values, which we refer to as the target parameters, ***θ*_*_**, corresponding to each Case considered. We take care to ensure that the three *in silico* experiments lead to typical *C*(*t*) data, as we demonstrate in Fig 2. The data from these experiments is treated as *observed* data, denoted **X**_obs_. Then, we use an ABC approach to explore, and quantify, how well the target values of ***θ*** can be estimated using the observed data. In particular, we are interested in the effect of varying the duration over which the observation data is collected, *T*.

In the absence of any experimental observations, information about ***θ*** is characterised by a specified prior distribution (Gelman et al., 2004, Sunnaker et al., 2013). For our choices of *α* and *β*, we set the prior to be
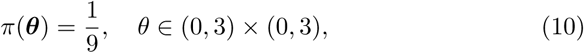

which is a uniform distribution across (*α*, *β*) ∈ (0, 3) × (0, 3).

We summarise data, **X**, with a lower-dimensional summary statistic, *S*. Under a Bayesian framework, the information from the prior is updated by the likelihood of the observations, *p*(*S*_obs_ |***θ***), to produce posterior distributions, *p*(***θ***|*S*_obs_), of ***θ***. In this study, we use the most fundamental ABC algorithm, known as ABC rejection (Liepe et al. 2014; Tanaka et al., 2006; Sunnaker et al., 2013). Our aim is to quantify the trade off between the duration of the experiment, *T*, and the reduction in uncertainty of the value of ***θ*** as well as the information gain.

In this work, we choose *S* to be the number of agents observed at equally spaced intervals of 24 hours. Let *N*_obs_(*t*) and *N*_sim_(*t*) denote the number of agents present in the observed data and a simulated cell proliferation assay at time t, respectively. We choose a discrepancy measure, *ρ*(*S*_obs_, *S*_sim_), to be the cumulative sum of the square difference between *N*_sim_(*t*) and *N*_obs_(*t*) at each 24 hour interval, up to the duration of the experiment, *T*, such that
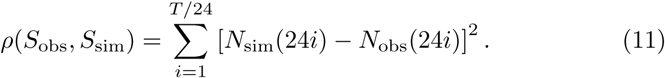

With these definitions, the ABC rejection algorithm is given by Algorithm 1.

#### Algorithm 1 ABC rejection sampling

1: Set *P_m_* = 0.579, *P_p_* = 0.002, *Δ* = 25 *μ*m, *τ* = 0:0417 hours, *N*(0) = 55.

2: Draw *θ_i_* ~ *π*(*θ*).

3: Simulate cell proliferation assay with *θ_i_*.

4: Record 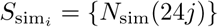; *j* = 1, 2, 3, 4.

5: Compute 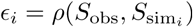, given in Eq 11.

6: Repeat steps 2-5 until 10^6^ samples 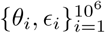 are simulated.

7: Retain a small proportion, *u* = 0.01, with the smallest discrepancy, *ϵ_i_*, as posterior samples.

To present and perform calculations with posterior distributions, we use a kernel density estimate with grid spacing 0.01 to form an approximate continuous posterior distribution from the samples. We do this using the ksdensity function in the MATLAB Statistics Toolbox (Mathworks, 2017). All ABC posterior results presented in the main paper correspond to retaining the 10,000 simulations out of 1,000,000 simulations with the smallest discrepancy, giving *u* = 0.01. To confirm that our results are insensitive to this choice of u we also present equivalent results with *u* = 0.02 in the Supplementary Material document.

#### 2.2.1 Kullback-Leibler divergence

To quantitatively compare posterior distributions, we calculate the Kullback-Leibler (KL) divergence (Kullback and Leibler 1951; Burnham and Anderson 2002), *D_KL_*(*p*||*π*), for each posterior distribution. The KL divergence is a measure of the information gain in moving from the prior, *π*(***θ***), to the posterior, *p*(***θ***| *S*_obs_), in Bayesian inference, and is defined as
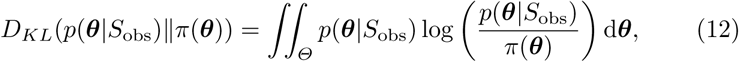

where *Θ* = (0, 3) × (0, 3) is the prior support. To calculate *D_KL_*(*p*(***θ***|*S*_obs_) ||*π* (***θ***)) we use quadrature to estimate the integral in Eq (12), taking care to ensure that the result is independent of the discretisation. Note that *D_KL_* is a measure of the amount of information gained when moving from the prior distribution to the posterior distribution.

#### 2.2.2 Other measures

We also make use of several other measures to help quantify various properties of the posterior densities. For each Case we always know, in advance, the target parameter values, ***θ***_*_, and we also estimate the mode, ***θ***_m_, using the kernel density estimate. Note that the mode is the value of ***θ*** corresponding to the maximum posterior density,
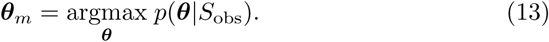

It is useful to report the posterior density at the target, *p*(***θ*_*_|***S*_obs_), for various values of *T*. It is also instructive to report the posterior density at the mode, *p*(***θ****m*|*S*_obs_), for various values of *T*. Another useful measure is the Euclidean distance between the target and the mode, given by
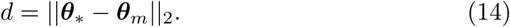

## 3 Results and Discussion

Results from a typical cell proliferation assay are shown in Fig 1(a)-(c). The cell density profile, shown in Fig 1(d), increases approximately linearly with time. This indicates that the experimental duration is not long enough for us to observe crowding effects, which occur at higher densities, and cause the net growth rate to reduce so that cell density profile, *C*(*t*), becomes concave down at later times. Therefore, by using typical experimental data, it is unclear whether the growth process follows a classical logistic model, or some other more general growth model.

To provide further insight into the limitations of this standard experimental design, we show results from the discrete model in Fig 2(a) for a standard experimental duration of *T* = 24 hours, for three different crowding functions. These results show several interesting features: (i) the cell density profile for each Case appears to increase linearly with time, which is similar to the experimental results in Fig 1(d); (ii) it is difficult to distinguish between the three different profiles, despite each profile corresponding to a different crowding function; and (iii) comparing the cell density profiles of a single realisation in Fig 2(a) to the expected behaviour in Fig 2(c) confirms that the expected cell density profiles for each Case are similar for the first 24 hours.

To examine when crowding effects begin to significantly influence the cell density profile, we perform simulations over longer durations of time. In particular, we examine *T* ≤ 96 hours. Results for a single realisation in Fig 2(b) show that the cell density profiles for each Case are indistinguishable during the first 24 hours. However the profile for each Case does become increasingly distinguishable at times greater than 24 hours. For example, each Case is clearly discernable by 72 hours. Comparing the cell density profiles of a single realisation in Fig 2(b) to the expected behaviour in Fig 2(d) confirms that each Case is only distinguishable at times greater than 24 hours. These observations motivate several questions that we will explore. The two main questions we focus on are: (i) what experimental duration is required to reliably distinguish between Cases 1, 2 and 3; and, (ii) can we quantify the trade off between allowing the experiment to run for a sufficiently long period of time to distinguish between the Cases, while still minimising the duration of the experiment.

To quantify the increase in information we can obtain by running the experiment for longer durations of time, we attempt to recover the parameters in the crowding function for each Case using ABC to produce a posterior distribution for *α* and *β*, which we refer to as the ordered pair (*α*, *β*). To achieve this aim, we produce *in silico* observed data, using a target parameter set for each Case: Case 1 corresponds to (*α, β*) = (1, 1); Case 2 corresponds to (*α*, *β*) = (2, 1); and Case 3 corresponds to (*α, β*) = (1, 2). All other parameters in the simulations are held fixed at the values given previously.

The data we use to perform inference takes the form of the size of the population, *N*(*t*), recorded at equally spaced intervals, each of duration 24 hours. In particular, we examine the effect of varying the total duration of the experiment, *T*. This means that if we consider an experimental design with *T* = 24 hours, then we record *N*(24) only. In contrast, if we consider an experimental design with *T* = 72 hours, we record *N*(24), *N*(48) and *N*(72). Overall, we examine four durations, *T* = 24, 48, 72 and 96 hours.

Results in Fig 3(a)-(d) show the bivariate posterior distributions of *α* and *β* for Case 1, with *T* = 24, 48, 72 and 96 hours, respectively. Recall that the target parameters for Case 1 are (*α, β*) = (1, 1). The results indicate that the choice of prior, π(θ), on the domain (0, 3) × (0, 3), is reasonable because the posterior distribution has full support within this region. The distribution in Fig 3(a) shows there are many parameter combinations that are likely to match the observed data, with *T* = 24 hours. This observation is consistent with the results in Fig 2(a) where we observe that setting *T* = 24 hours is insufficient to distinguish between the three Cases. Comparing the posterior distributions in Fig 3(a)-(d), we see that increasing *T* leads to a narrowing of the posterior distribution, and the mode of the distribution moves toward the target parameter combination. For this Case, we see the largest benefit when increasing *T* from 48 to 72 hours. For example, for *T* = 48 hours, the mode of the distribution is (1.82, 2.16), which means that each parameter estimate is almost double each target value. In contrast, the mode of the distribution at *T* = 72 hours is (1.06, 0.95), so each parameter is able to be estimated within 6% of the target.

**Fig. 3.**
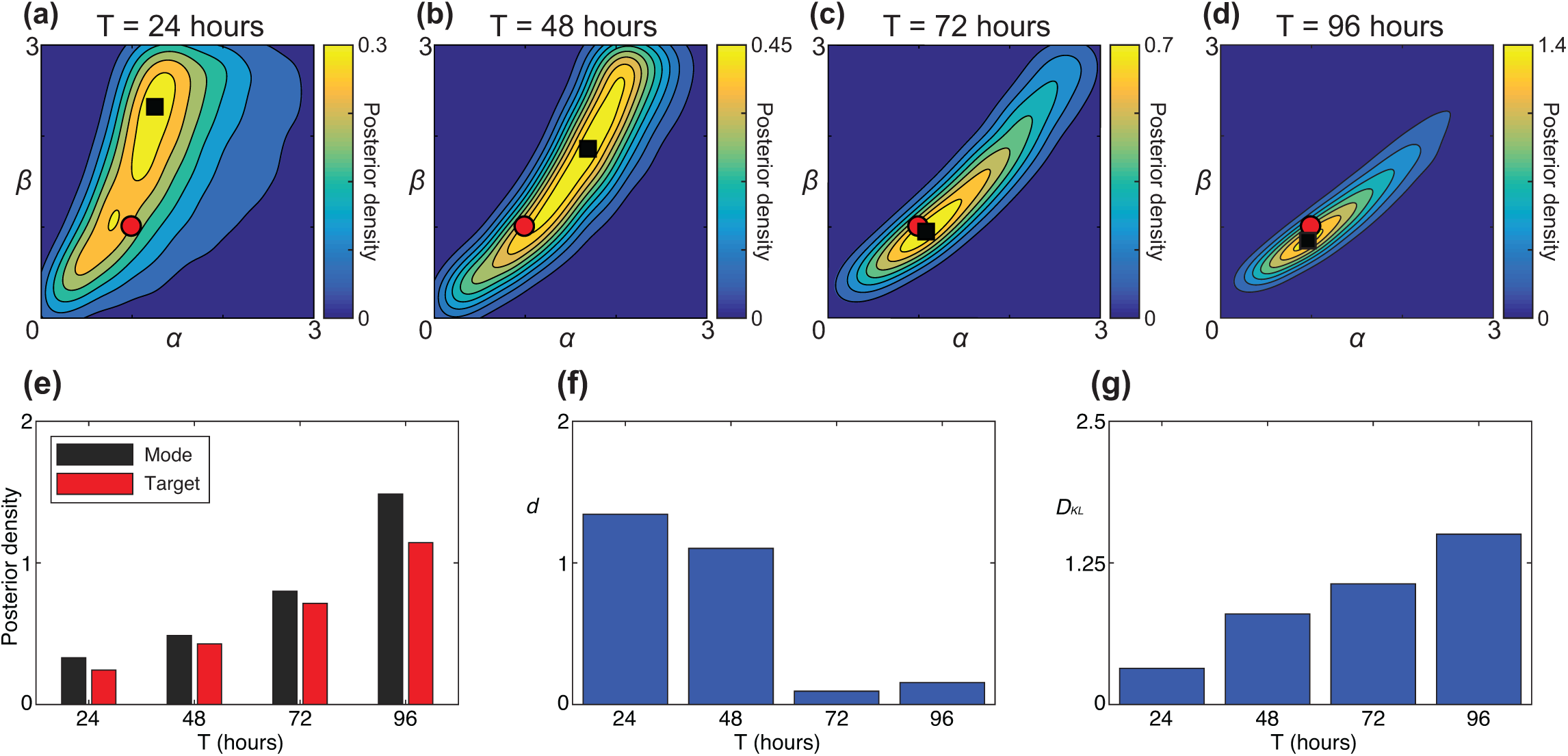
Posterior distributions for Case 1: (*α, β*) = (1,1). (a)-(d) ABC posterior distributions for: (a) *T* = 24 hours; (b) *T* = 48 hours; (c) *T* = 72 hours and (d) *T* = 96 hours. The posterior distributions are approximated using the best 10,000 samples from 1,000,000 prior samples (*u* = 0.01), as measured by *ρ,* given by Eq 11. The red circles show the location of the target parameters used to generate the observed data (*α* = 1, *β* = 1). The black squares indicate the mode of the posterior distribution. The modes are (1.32, 2.43), (1.82, 2.16), (1.06, 0.95) and (0.96, 0.86) in (a)-(d), respectively. (e)-(g) Show measures of accuracy and precision. (e) Quantitatively compares the posterior density at the mode and the target parameter values. (f) Shows *d,* the Euclidean distance between the mode and target parameter values, given by Eq 14. (g) Shows *D_KL_,* the Kullback-Leibler divergence from the prior, for each posterior distribution, given by Eq 12.

To quantify the properties in the posterior distributions, Fig 3(a)-(d), there are many features that we may consider. Figure 3(e) compares the posterior density at the target parameter values and the maximum posterior density of the distribution, which corresponds to the mode. The maximum posterior density increases with *T*, confirming that the posterior distribution narrows as the duration of the experiment is increased. Results in Figure 3(f) show that d eventually decreases with *T*, indicating that the mode of the distribution moves towards the target as *T* increases. Together, these results show that the density at the mode is close to the density at the target, and that both these quantities increase with *T*. This indicates that the target parameter combination is always as likely as the mode. Results in Fig 3(g) shows how the KL divergence (Eq 12) also increases with *T*. We see that the largest gain in information for this Case occurs when *T* is increased from 24 hours (*D_KL_* = 0.33) to 48 hours (*D_KL_* = 0.84). The quantitative measures in Fig 3(e)-(g) suggest that there is always value in increasing *T*, however the value of increasing *T* varies. For example, there is a substantial benefit in extending the experiment from *T* = 48 to 72 hours, whereas the benefit in extending the experiment from *T* = 72 to 96 hours is less pronounced.

Results in Fig 4(a)-(d) and Fig 5(a)-(d) show the bivariate posterior distributions of *α* and *β* for Cases 2 and 3, respectively. Note that all data presented for Cases 2 and 3 is given in the same format as used for the results corresponding to Case 1 in Fig 3. As before, we always observe a narrowing of the posterior distribution as T increases. Results in Fig 4(e) and Fig 5(e) clearly show that the target parameter combination becomes more likely as *T* is increased. Data for *d* in Fig 4(f) confirms that the distance between the target and the mode is reduced for larger values of *T*. Data for d in Fig 5(f) shows that the distance between the target and the mode increases, at first, when *T* is increased from 24 to 48 hours. However, the most important feature is that *d* always decreases eventually for large enough *T*. Again, as *T* is increased, *D_KL_* increases in both Fig 4(g) and Fig 5(g).

**Fig. 4.**
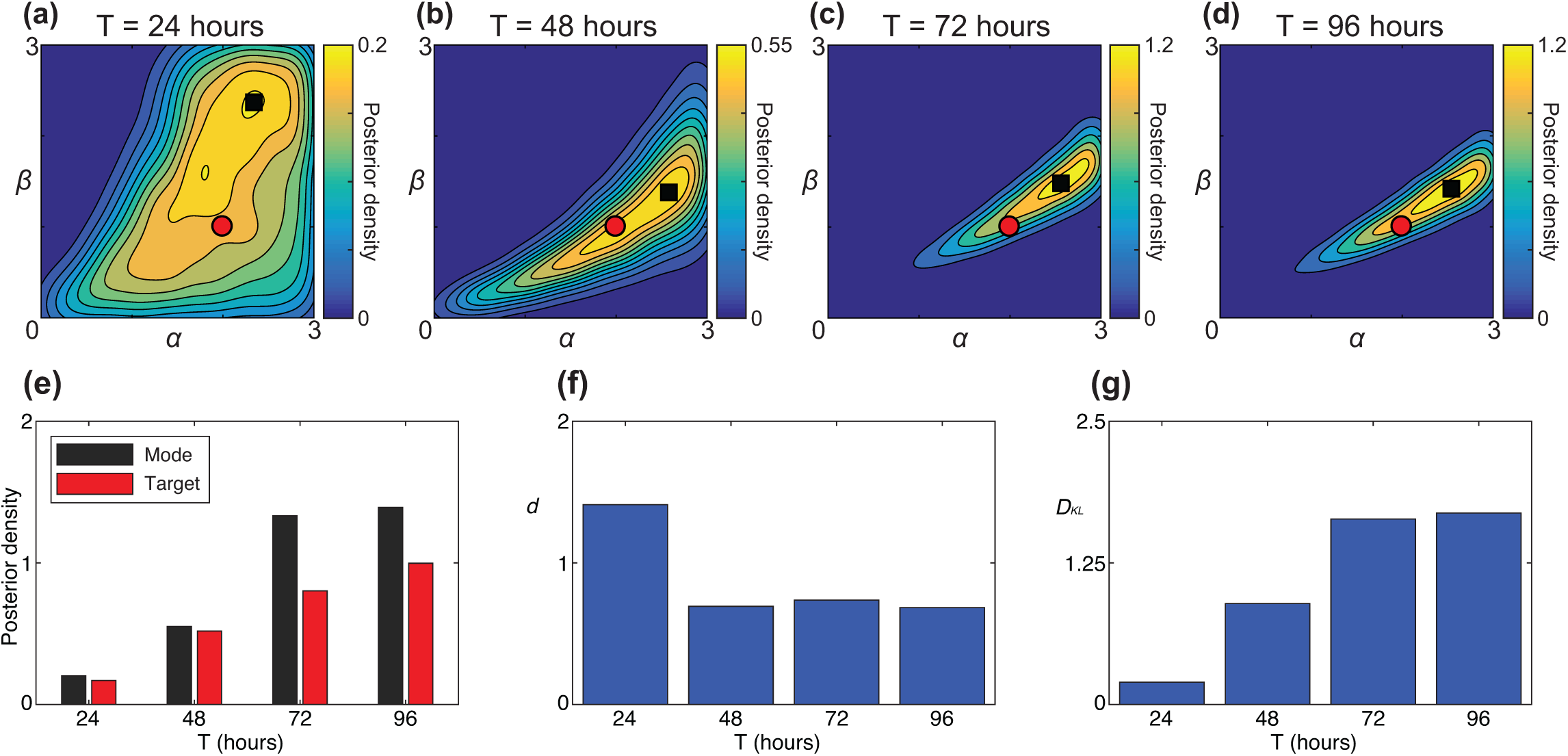
Posterior distributions for Case 2: (*α, β*) = (2,1). (a)-(d) ABC posterior distributions for: (a) *T* = 24 hours; (b) *T* = 48 hours; (c) *T* = 72 hours and (d) *T* = 96 hours. The posterior distributions are approximated using the best 10,000 samples from 1,000,000 prior samples (*u* = 0.01), as measured by *ρ,* given by Eq 11. The red circles show the location of the target parameters used to generate the observed data (*α* = 2, *β* = 1). The black squares indicate the mode of the posterior distribution. The modes are (1.89, 1.81), (2.55, 1.38), (2.54, 1.46) and (2.53, 1.41) in (a)-(d), respectively. (e)-(g) Show measures of accuracy and precision. (e) Quantitatively compares the posterior density at the mode and the target parameter values. (f) Shows *d,* the Euclidean distance between the mode and target parameter values, given by Eq 14. (g) Shows *D_KL_,* the Kullback-Leibler divergence from the prior, for each posterior distribution, given by Eq 12.

**Fig. 5.**
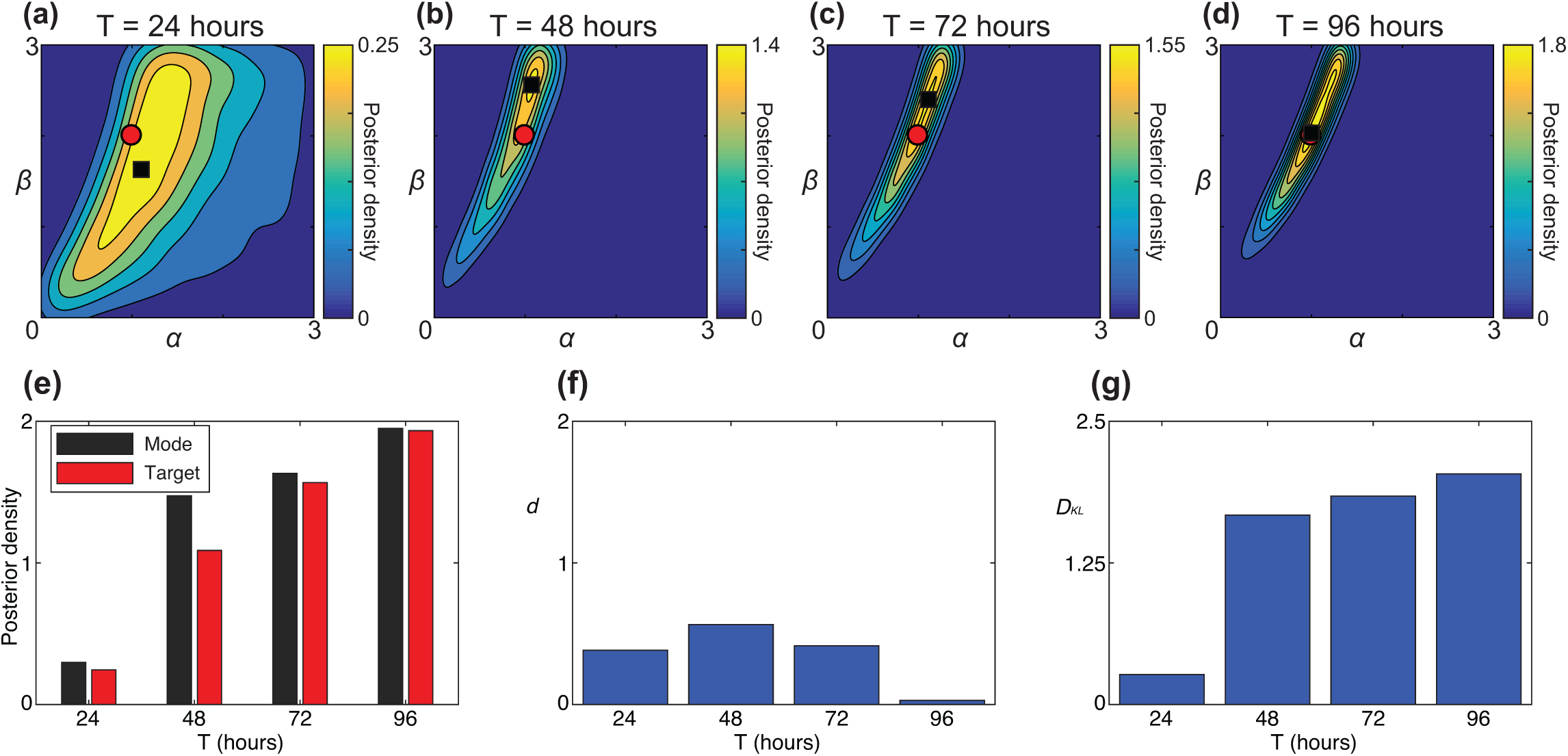
Posterior distributions for Case 3: (*α*, *β*) = (1, 2). (a)-(d) ABC posterior distributions for: (a) *T* = 24 hours; (b) *T* = 48 hours; (c) *T* = 72 hours and (d) *T* = 96 hours. The posterior distributions are approximated using the best 10,000 samples from 1,000,000 prior samples (*u* = 0.01), as measured by *ρ,* given by Eq 11. The red circles show the location of the target parameters used to generate the observed data (*α* = 1, *β* = 2). The black squares indicate the mode of the posterior distribution. The modes are (1.13, 1.70), (1.09, 2.57), (1.20, 2.67) and (1.03, 2.11) in (a)-(d), respectively. (e)-(g) Show measures of accuracy and precision. (e) Quantitatively compares the posterior density at the mode and the target parameter values. (f) Shows *d,* the Euclidean distance between the mode and target parameter values, given by Eq 14. (g) Shows *D_KL_,* the Kullback-Leibler divergence from the prior, for each posterior distribution, given by Eq 12.

Overall, the essential trends in Fig 4 and Fig 5 are consistent with those in Fig 3, namely: (i) the standard choice of *T* = 24 hours is insufficient to determine the parameters in the crowding function and hence it is impossible to reliably distinguish between classical logistic growth and more general logistic growth models; and, (ii) as the value of *T* is increased, our ability to recover the parameters in the crowding function increases. However, certain details differ between the cases. For example, choosing *T* = 72 hours allows us to recover estimates of *α* and *β* to an accuracy of at least 6, 46 and 34% in Cases 1, 2 and 3, respectively. Therefore, with this choice of *T* we are able to recover the parameters for Case 1 relatively accurately. In contrast, if we choose *T* = 96 hours, we recover estimates of α and *β* to an accuracy of at least 14, 41 and 6% in Cases 1, 2 and 3, respectively. Therefore, with this choice of *T* we are able to recover the parameters for Cases 1 and 3 relatively accurately, yet Case 2 remains relatively unclear.

## 4 Conclusion

In this work, we implement a random walk model to simulate a cell proliferation assay. In particular, we focus on exploring whether the typical experimental design of a cell proliferation assay, with *C*(0) ≈ 0.1, **λ** ≈ 0.05 /hour and *T* = 24 hours, is sufficient to make a clear distinction between classical logistic growth and more general logistic growth models. We are motivated to explore this question because many theoretical modelling studies choose to represent cell proliferation with the classical logistic model, yet this assumption is rarely tested using experimental data. Furthermore, there is a growing awareness in the mathematical biology literature that the choice of using a classical logistic model can be inappropriate. For example, Sarapata and de Pillis (2014) show that a range of tumour growth data is more accurately predicted using a generalised logistic model rather than the classical logistic model. Therefore, the question of whether standard designs of cell proliferation assays can make a clear and unambiguous distinction between classical logistic growth and more general logistic growth is important as cell proliferation assays are commonly employed. It is currently unclear whether the standard experimental design is sufficient to distinguish between different sigmoid growth mechanisms. This study is the first time that a stochastic individual based model has been used to explore the optimal duration of a cell proliferation assay. In particular, we explore how to choose the duration of the assay to reliably distinguish between different types of growth models.

One of the main conclusions of our study is that the typical experimental design for a cell proliferation assay, with *C*(0) ≈ 0.1, **λ** ≈ 0.05 /hour and *T* = 24 hours, can not be used to make a distinction between classical logistic growth and more general logistic growth. Further, we use our stochastic modelling and parameter inference tools to explore how the experimental design can be altered so that this distinction can be made with confidence. In particular we explore the option of increasing the duration of the experiment, *T*. Our parameter inference results show that increasing *T* always provides more information about the crowding function parameters. However, the trends are subtle, and there is no simple guideline for prescribing the ideal experimental duration that one could implement in practice. Our results show that we can recover the crowding function for the case of classical logistic growth (Case 1: *α* = 1, *β* = 1) to within an accuracy of 6% if the experimental duration is increased to *T* = 72 hours. Beyond this duration, we encounter diminishing returns for this Case. For example, further increasing the duration of the experiment to *T* = 96 hours leads to only a small increase in additional information about the crowding function. In other cases where we consider generalised logistic growth (Case 2: *α* = 1, *β* = 2), we see that the parameter estimates remain relatively poor, even if the experimental duration is increased to *T* = 72 hours. For Case 2 we recover the parameters to an accuracy of within 33% if *T* = 72 hours, and to within 5% if *T* = 96 hours. Therefore, it is not possible to make a simple conclusion that cell proliferation assays ought to be conducted until *T* = 48 or *T* = 72 hours since the increase in information with *T* is subtle. Despite this complication, our results certainly show that the standard choice of *T* = 24 hours is insufficient, and that the experiment ought to be conducted for a long as practically possible.

One aspect of a cell proliferation assay that we have not explored is the dependence of the results on the initial cell density, *C*(0). All results in this work, both the *in vitro* experimental data in Fig 1, and the *in silico* data in Figs 2-5, deal with initial densities of *C*(0) ≈ 0.1, where *C* = 1 corresponds to the maximum carrying capacity of the confluent monolayer. This initial density corresponds to a fairly standard choice of initiating a cell proliferation assay with approximately 20,000 cells placed into the wells of a 24-well tissue culture plate where each well has a diameter of approximately 15 mm. Alternatively, a similar initial density can be obtained by initiating a cell proliferation assay with approximately 10,000 cells placed into the wells of a 96-well tissue culture plate, where each well has a diameter of approximately 9 mm. While it is true that crowding effects in a cell proliferation assay might be more clearly discernable by initiating the experiment with larger numbers of cells, we warn against this for two reasons. First, from a practical point of view, our experience in initiating a two-dimensional *in vitro* cell biology assay with large numbers of cells is problematic as the cells can tend to cluster together, and pile up in the vertical direction instead of spreading as a monolayer (Treloar et al. 2013). Second, established methods for initiating cell proliferation assays with *C*(0) ≈ 0.1 are perfectly well suited to observe the low density exponential phase of the growth process, which is important to estimate the intrinsic proliferation rate, **λ**. For example, the data shown in Fig 1(a)-(c) corresponds to a cell proliferation assay initialised with 20,000 cells in a 24-well tissue culture plate, and results in Fig 1(d) show *C*(*t*) grows linearly over the first 24 hours. This result is consistent with the early part of the growth process where we expect 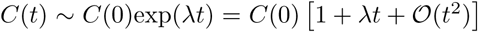. Therefore, we do not suggest that the standard experimental design for a cell proliferation assay ought to be altered by increasing *C*(0). This is why, throughout this study, we have treated *λ* and *C*(0) as known, constant values, in the experimental design.

All of the results presented here have focused on exploring whether we can make a reliable distinction between classical logistic growth and more general logistic growth in a cell proliferation assay. To achieve this we use *in silico* simulations in which the crowding function can be specified. While the discrete simulation algorithm can be used to model a cell proliferation assay with any crowding function, *f*(*C*), to illustrate the key points of our study we focus on three particular cases. Case 1 corresponds to classical logistic growth, while Cases 2 and 3 are examples of more general logistic growth. Of course, the methods outlined in this work apply equally well to any other choice of crowding function. Furthermore, while all crowding functions explored here involve two parameters, *α* and *β*, it is possible that other choices of crowding function might contain additional parameters. Under these conditions, the procedures described here to quantitatively measure the potential for parameter recovery as a function of the experimental design apply in exactly the same way regardless of the number of unknown parameters in the crowding function.

## 5 Acknowledgments

This work is supported by the Australian Research Council (DP140100249, DP170100474). Computational resources were provided by the High Performance Computing and Research Support Group. We thank the two anonymous referees for their helpful comments.

